# Chondroitin sulfate degradation bolsters *Proteus mirabilis* growth and colonization of the catheterized urinary tract

**DOI:** 10.64898/2026.03.02.708568

**Authors:** Benjamin C. Hunt, Vitus Brix, Namrata Deka, Brian S. Learman, Aimee L. Brauer, Brianna M. Shipman, Nicole De Nisco, Chelsie E. Armbruster

## Abstract

Glycosaminoglycans (GAGs) are negatively charged polysaccharides composed of repeating disaccharide units and are essential components of the extracellular matrix throughout numerous tissues. The bladder urothelium has a thick protective GAG layer that primarily consists of chondroitin sulfate (CS), heparan sulfate (HS), and hyaluronic acid (HA), and urinary tract pathogens must either degrade or otherwise circumvent this layer to infect the urothelium. In this study, we investigated GAG degradation by *Proteus mirabilis,* a common and persistent colonizer of the catheterized urinary tract. Almost all *P. mirabilis* urinary tract isolates harbor a putative chondroitin endolyase (PMI2127), exolyase (PMI2128), and sulfatase (PMI2124). By generating mutant and complemented strains of these genes, we determined that *P. mirabilis* strain HI4320 degrades multiple forms of CS under numerous culture conditions, including during growth in human urine, and can use CS degradation products as a carbon source. Sulfatase and endolyase activity were required for efficient degradation of all CS types, while the exolyase only contributed to using CS-B or CS-C as carbon source. Interestingly, only endolyase activity contributed to colonization in a murine model of CAUTI, although the colonization defect was even more pronounced when the endolyase and exolyase were both disrupted. The colonization defect was specific to the CAUTI model, likely due to the impact of catheterization on the GAG landscape of the bladder. Limiting CS degradation by *P. mirabilis* may therefore represent a strategy for reducing risk of ascending infection in catheterized patients.

**Importance:** Glycosaminoglycans (GAGs) are a family of negatively charged heteropolysaccharides that are ubiquitously expressed throughout the body, forming a significant component of the extracellular matrix and a luminal GAG layer in the bladder. This GAG layer functions as a physical barrier for the bladder surface, protecting it from bacterial infection. Disruption of this barrier through physical forces, such as catheter insertion, or enzymatic degradation by bacteria may contribute to infection outcomes. In this study, we defined the contribution of three putative chondroitin sulfate degrading enzymes (PMI2124, PMI2127, PMI2128) to the pathogenesis of a common pathogen in the catharized urinary tract, *Proteus mirabilis.* We found that *P. mirabilis* can utilize chondroitin sulfate as a carbon source, and that chondroitin sulfate degradation contributes to infection in a model of catheterized urinary tract infection. This work contributes to a growing understanding of how uropathogens subvert host defenses and acquire nutrients within the bladder.

## Introduction

The urinary bladder relies on a highly specialized epithelial barrier known as the urothelium to maintain normal function despite constant exposure to urine, a complex fluid containing metabolic waste products, ions, and microorganisms (1–3). The urothelium is divided into three components: the apical, lateral, and basal cell layers. The apical layer consists of terminally-differentiated umbrella cells that express uroplakins, hexagonal crystalline membrane proteins that contribute to the flexible permeability barrier of the bladder, as well as a luminal glycocalyx of glycosaminoglycans (GAGs) (2, 4, 5). GAGs are negatively charged heteropolysaccharides consisting of repeating disaccharide units of an *N*-acetylated hexosamine and uronic acid, and they are categorized into four major classes based on the type of hexosamine and uronic acid they contain and the glycosidic linkage between these units (5–11). In the bladder, the GAG layer is a mucus-like coating composed primarily of chondroitin sulfate (CS), along with heparan sulfate (HS) and hyaluronic acid (HA) (5–7).

The bladder GAG layer functions as a physical shield for the urothelium, limiting epithelial permeability, repelling urinary solutes, and inhibiting bacterial adherence (3, 5, 7, 12). Disruption of the GAG layer has been increasingly recognized as a key contributor to bladder inflammation, pain, and susceptibility to infection (5, 9, 13). Alterations in the composition or function of this GAG barrier have been implicated in multiple bladder disorders, such as interstitial cystitis/bladder pain syndrome (IC/BPS), where increased epithelial permeability is thought to drive chronic pain, urgency, and frequency (14). With respect to infection, we recently demonstrated that GAG degradation by *Enterococcus faecalis* is an important contributor to colonization and tissue dissemination during catheter associated urinary tract infection (CAUTI) and bacteremia (15). Nguyen et. al 2022 also demonstrated that GAG degradation is an activity diverse bacterial species participate in, including several species known to be associated with the urinary tract microbiota or urinary tract pathogens (15, 16). Considering its central importance in urothelial defenses, it is not surprising that multiple uropathogenic bacteria produce enzymes capable of degrading GAGs (15, 17–19). Enzymatic cleavage of GAGs may weaken the urothelial barrier, increase epithelial permeability, and facilitate bacterial adherence and persistence (17, 20, 21). Thus, GAG degradation by uropathogens may be a contributor to infection, barrier dysfunction, and more severe disease (21–23).

Despite growing recognition of the importance of the urothelial GAG layer in bladder health, the contribution of bacterial GAG degradation to UTI and CAUTI represents a gap in knowledge that is just beginning to be explored. One uropathogen that has been reported to degrade GAGs is *P. mirabilis*, a Gram-negative bacterium that is one of the leading causes of CAUTI (24–28). *P. mirabilis* has numerous virulence factors that promote colonization and persistence in the catheterized urinary tract, including adhesins, flexible metabolic pathways for optimal growth when using amino acids and peptides as primary nutrients, and potent urease activity for nitrogen acquisition coupled with crystalline biofilm formation. Since GAG degradation releases disaccharides such as *N*-acetylglucosamine (GlcNAc) and *N*-acetylgalactosamine (GalNAc) (5–11), GAG degradation may represent another factor employed by this adaptable uropathogen to facilitate optimal growth, colonization, tissue invasion (25, 29).

*P. mirabilis* has two predicted chondroitin sulfate degradation enzymes, PMI2127 and PMI2128. PMI2127 is predicted to be a chondroitin ABC endolyase, which cleaves the internal bond between GlcA and GalNAc, while PMI2128 is predicted to be a chondroitin ABC exolyase, which cleaves off disaccharide units from the non-reducing ends of the sugar chains. *P. mirabilis* also encodes a predicted exported sulfatase, PMI2124. In this study, we defined the contribution of these three gene products to *P. mirabilis* GAG degradation, growth *in vitro,* and pathogenesis in three different infection models. We show that *P. mirabilis* readily degrades multiple types of CS but not HS or HA. We further show that PMI2124 and PMI2127 are required for degradation of all CS types, while PMI2128 is only required for specific CS subsets. Additionally, CS degradation was important for optimal colonization of the urinary tract in a murine CAUTI model, but not in a model of ascending UTI without a catheter. Our data therefore indicates that CS degradation is specifically important to *P. mirabilis* infection in the catheterized bladder environment, and paves the way for further examining the impact of catheterization on the GAG landscape of the urinary tract.

## Results

### *P. mirabilis* HI4320 has two functional chondroitinases

*P. mirabilis* strain HI4320 has two predicted chondroitinases: an endolyase (PMI2127) and an exolyase (PMI2128), as well as a putative sulfatase, PMI2124, that may play a role in chondroitin degradation (Figure 1A). Analysis of 1,027 *P. mirabilis* genomes isolated from urine revealed near-universal presence of the three GAG genes: PMI2124 (99.4%), PMI2127 (99.4%), and PMI2128 (99.2%) (Figure 1B). All three genes co-occurred in 1,019 genomes (99.2%) with high sequence conservation (86.4-100% identity), including in 27 recent isolates from catheterized nursing home residents (30).

**Figure 1.**
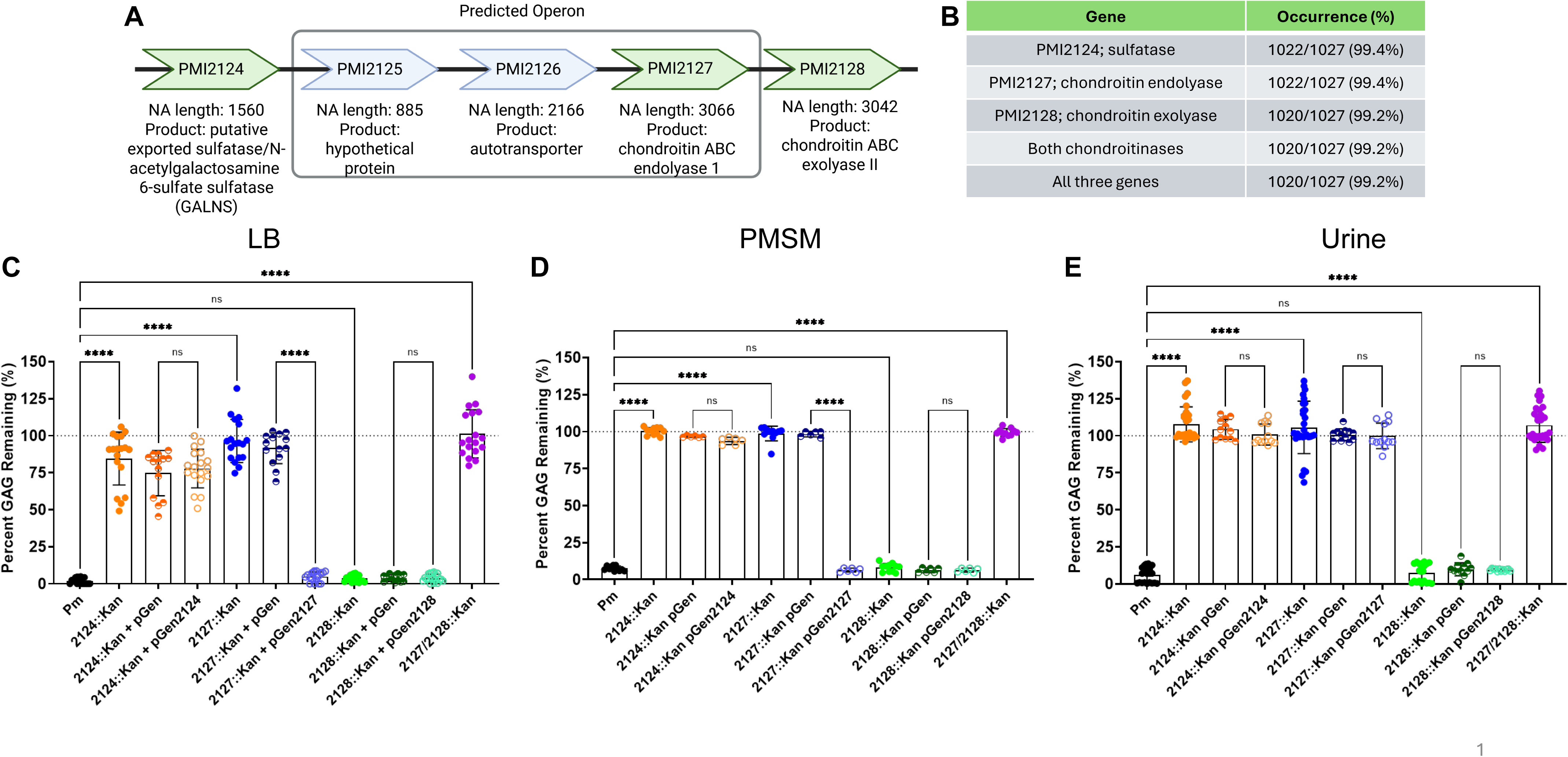
*Proteus mirabilis* chondroitin sulfate degradation genes and activity in different media. A) Predicted sulfatase, chondroitin ABC endolase, and chondroitin ABC exolyase (green), as well as the operon organization in *P. mirabilis* strain HI4320, as predicted by Bio-Cyc. B) Analysis of 1,027 *P. mirabilis* genomes for the presence or absence of PMI2124, PMI2127, and PMI2128 with ≥80% identity and ≥80% query coverage. (C-E) Percent chondroitin sulfate remaining after 48hrs of growth of *P. mirabilis* HI4320 or indicated mutant, empty vector, and complemented strains in LB media (C), *Proteus* minimal salts media (PMSM, D), or human urine (E). Data represent mean ± SD of at least three independent experiments. Statistical comparisons represent One-way ANOVA with multiple comparison correction to wild-type Pm, **** p<0.0001.

To begin investigating the ability of *P. mirabilis* to degrade chondroitin sulfate (CS) and potentially other GAGs, we utilized the semi-quantitative GAG degradation assay developed by Nguyen et. al 2022. This assay relies on binding of high molecular weight GAGs to positively-charged bovine serum albumin at neutral pH, then inducing development of insoluble GAG-BSA complexes by acidifying the media with acetic acid. We generated three single mutants wherein PMI2124, PMI2127 or PMI2128 were knocked out, as well as a PMI2127/2128 double mutant to investigate if there is functional redundancy between the two chondroitinases. We first examined degradation of CS in rich laboratory media (LB supplemented with 2.5 mg/mL CS) and quantified the percentage of high molecular weight CS remaining after 48 hours. Wild-type *P. mirabilis* readily degraded all available CS, while disruption of either PMI2124 or PMI2127 abrogated degradation (Figure 1C). Disruption of PMI2128 seemed to have no effect on the amount of GAG degradation, suggesting that endolyase activity is primarily responsible for CS degradation under these conditions. Each mutant was complemented in trans by expressing the wild-type gene under its native promoter on a plasmid (pGEN). Complementation of PMI2127 fully restored chondroitin degradation, but PMI2124 did not. Lack of complementation could be due to polar effects of the mutation or to lack of sufficient expression from the native promotor. To investigate, we performed RT-qPCR analysis of strains grown in LB with or without 2.5 mg/mL of chondroitin sulfate mix (Supplemental Figure 1). Expression of PMI2124 was fairly constant over time in wild-type *P. mirabilis,* did not appear to be induced by the presence of CS, and remained constant in each of the mutant and complemented strains. In contrast, PMI2127 and PMI2128 were highly induced by CS at 18-24hrs of growth in some experiments but not all, and complementation enhanced expression by 20-80 fold wild-type levels. Importantly, disruption of PMI2124 did not substantially decrease expression of PMI2127 or PMI2128, suggesting that lack of activity in the PMI2124 mutant is not due to polar effects on the downstream CS lyases.

To examine the GAG specificity of PMI214 and PMI2127, the degradation assay was repeated in LB supplemented with either hyaluronic acid (HA) or heparan sulfate (HS). Wild-type *P. mirabilis* was unable to degrade either additional GAG substrate, and the mutants all exhibited comparable profiles to wild-type (Supplemental Figure 2). Thus, *P. mirabilis* specifically degrades CS through the action of a sulfatase and an endolyase under these experimental conditions.

We next investigated CS degradation under nutrient limited conditions, including in a minimal growth media (PMSM) and in human urine. Wild-type *P. mirabilis* and all mutants exhibited the same degradation profile in PMSM as in LB, and complementation of PMI2127 restored degradation while complementation of PMI2124 did not (Figure 1D). In human urine wild-type *P. mirabilis* again readily degraded CS, disruption of either PMI2124 or PMI2127 abrogated degradation, and loss of PMI2128 had no impact (Figure 1E). However, complementation of PMI2127 in human urine did not restore degradation, suggesting that regulation may be altered under these conditions.

Given the different nutrient environment of LB and human urine, we were curious whether the kinetics of CS degradation differ between media types. To this end, we measured CS degradation across a time course (2, 4, 6, 12, 18, 20, and 24 hours) in LB and urine. Degradation of CS began earlier during growth in urine (12 and 18 hours) than during growth in LB (20 hours, Figure 2). Thus, environmental conditions such as nutrient availability likely contribute to regulating the expression or activity of *P. mirabilis* chondroitinases.

**Figure 2.**
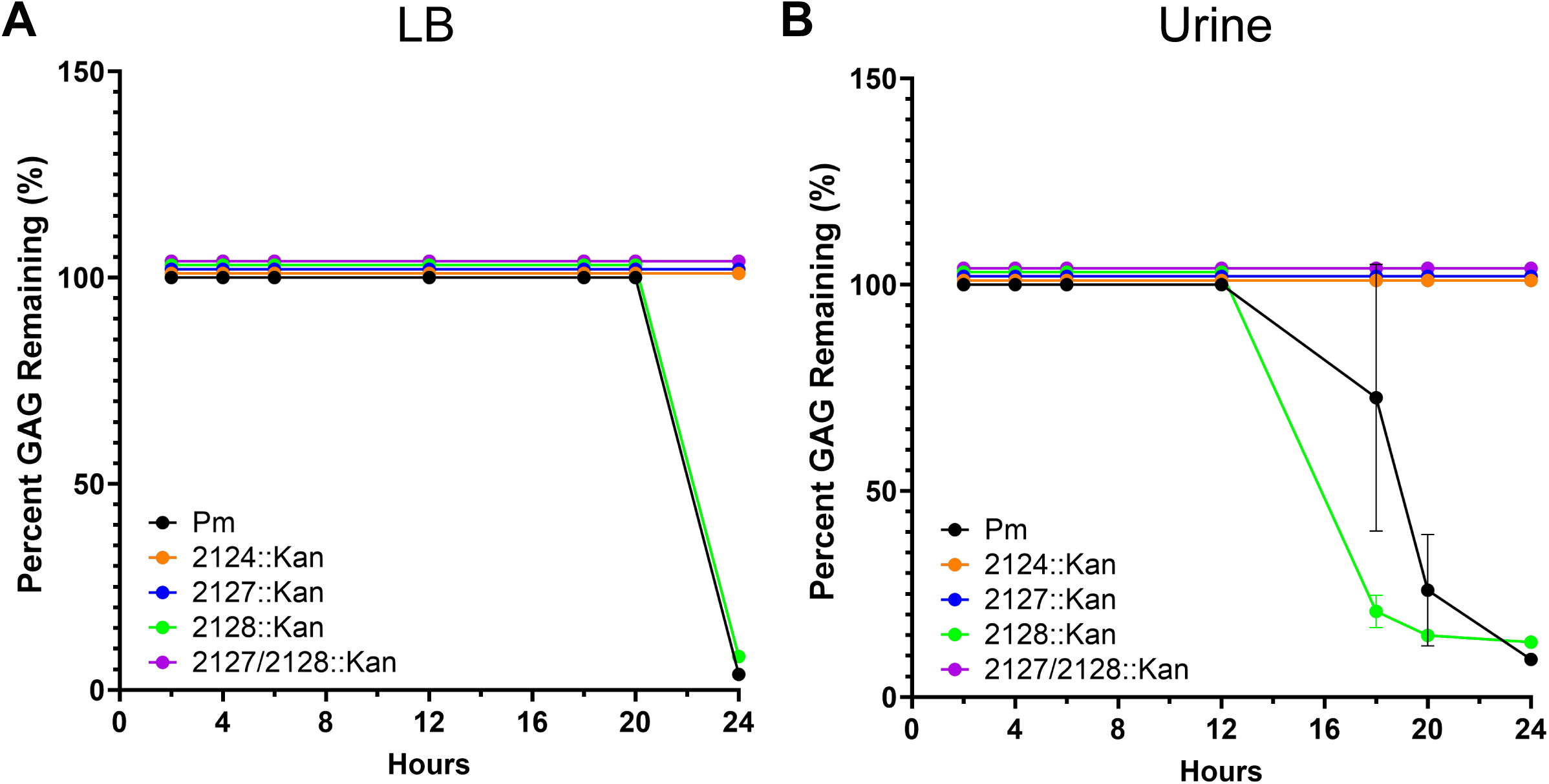
The timing of chondroitin sulfate degradation is influenced by media composition. Percent chondroitin sulfate remaining after 2, 4, 6, 12, 18, 20, and 24 hrs of growth of *P. mirabilis* HI4320 or indicated mutant strains in LB (A) or human urine (B). Data represent mean ± SD of at least three independent experiments.

CS is classified into different types based upon the sulfonation patterns present on the glucuronic acid – N-acetyl galactosamine backbone (10). Thus, to examine the specificity of *P. mirabilis* CS degradation, GAG degradation assays were repeated with chondroitin A (GlcA-GalNAc(4S)), B (GlcA(2S)-GalNAc(4s), or C (GlcA-GalNac(6S)). Degradation of the individual CS types mirrored the results observed for the unspecified CS mix in Figure 1: wild-type *P. mirabilis* readily degraded CS-A, CS-B and CS-C in each of the media conditions (LB, PMSM, urine), disruption of either PMI2124 or PMI2127 abrogated degradation, and complementation of PMI2127 restored degradation in LB and PMSM but not urine. (Figure 3). Overall, these data show that *P. mirabilis* is able degrade CS with different sulfonation patterns, including within physiologically relevant media such as human urine, and that PMI2124 and PMI2127 are essential for CS degradation under these conditions.

**Figure 3.**
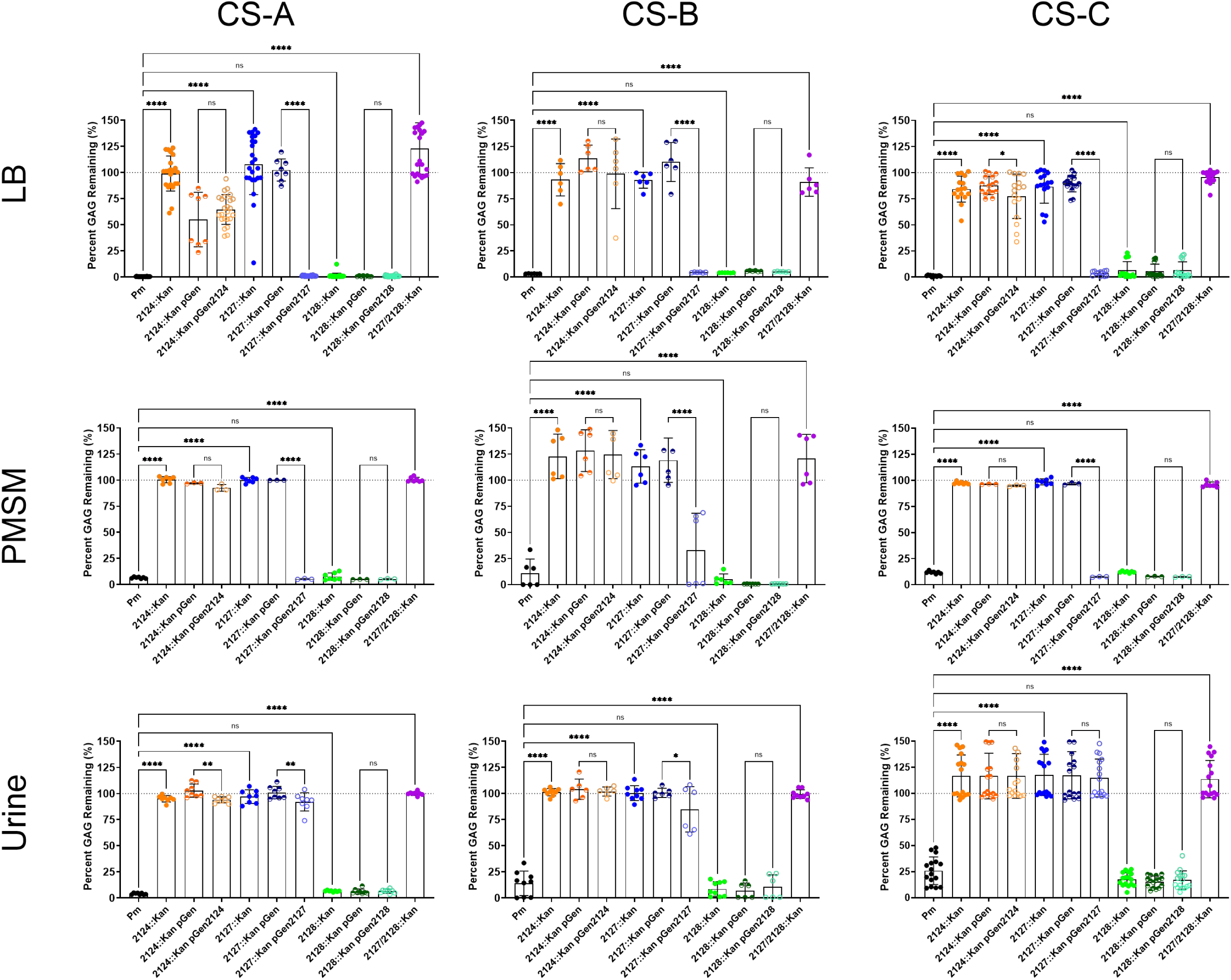
*P. mirabilis* can degrade multiple chondroitin sulfate types (CS-A, CS-B, and CS-C) in multiple media types. Graphs display the percent of each CS type (A, B, or C) remaining after 48 hrs growth of *P. mirabilis* HI4320 or indicated mutant, empty vector, and complemented strains in LB (top), PMSM (middle), or human urine (bottom). Data represent mean ± SD of at least three independent experiments. Statistical comparisons represent One-way ANOVA with multiple comparison correction to wild-type Pm condition, **** p<0.0001.

### *P. mirabilis* can utilize chondroitin sulfate as a sole carbon source for growth

Given that *P. mirabilis* is able to degrade CS, and that the GlcA and GalNAc residues that comprise CS can be utilized by various bacteria for growth (18, 31, 32), we next investigated whether *P. mirabilis* was able to utilize CS degradation products as a carbon source for growth in nutrient limiting conditions. Growth of *P. mirabilis* was assessed for 48 hours in PMSM with either no carbon source, 2.5 mg/mL glucose, 2.5 mg/mL of CS mixture, or 2.5 mg/mL of each individual CS type (Supplemental Figure 3 and Figure 4). Growth of wild-type *P. mirabilis* was less robust in PMSM with CS than glucose (maximum OD_600_ of ∼0.4 vs 0.6), but strain HI4320 was able to use CS as a sole carbon source. Interestingly, the growth pattern was seemingly diauxic, suggesting that *P. mirabilis* may preferentially degrade CS with certain sulfonation patterns first before switching to others (Supplemental Figure 3A and Figure 4B). A higher optical density was also achieved by wild-type when growing on the CS mixture than any individual CS type, providing further support for a potential sulfonation pattern preference.

**Figure 4.**
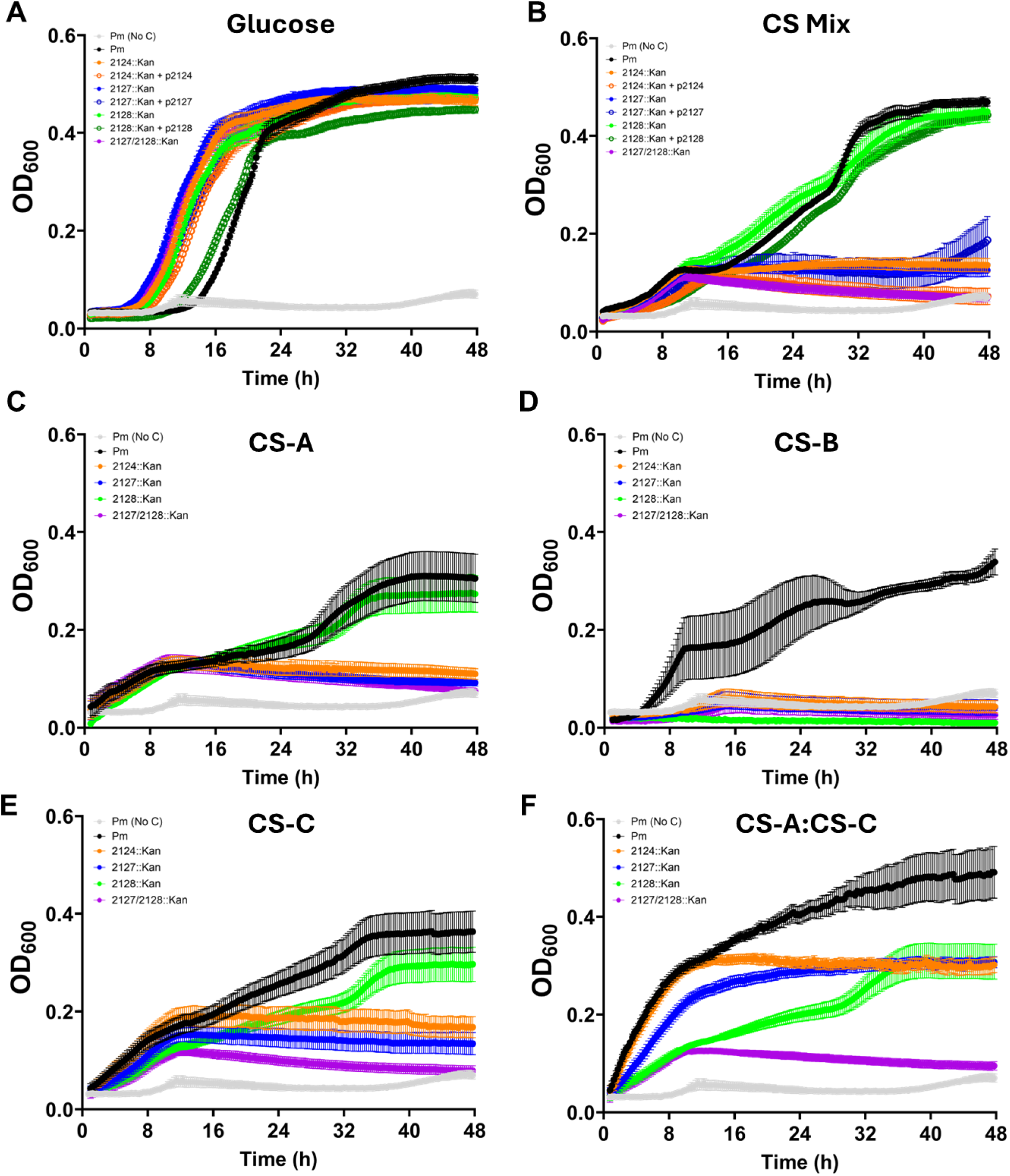
*P. mirabilis* growth with chondroitin sulfate as a sole carbon source. *P. mirabilis* HI4320 or indicated mutant, empty vector, and complemented strains were grown for 48hr in PMSM with A) 2.5 mg/mL glucose (gluc), B) chondroitin sulfate mix (CS Mix), C) chondroitin sulfate A (CS-A), D) chondroitin sulfate B (CS-B), E) chondroitin sulfate C (CS-C), or F) a 1:1 mixture of CS-A and CS-C. Growth was measured by OD600 readings every 15 minutes. Data represent mean ± SEM of at least three independent experiments with at least three replicates each.

Disrupting genes involved in CS degradation had no impact on growth with glucose as the carbon source (Figure 4A). In agreement with our degradation assay results, the ability to use any CS type as a sole carbon source was largely dependent on desulfatase and endolyase activity, as minimal growth was observed when either PMI2124 or PMI2127 were disrupted (Figure 4B). Complementation of PMI2127 resulted in a slight increase in growth on the CS mixture after 40 hours, suggesting at least partial restoration under this condition but not any of the individual CS types (Figure 4B), while complementation of PMI2124 had no effect. Interestingly, all mutants struggled to grow when supplied with CS-B as the sole carbon source, but PMI2128 failed to exhibit any growth under this condition (Supplemental Figure 3D and Figure 4D). Thus, exolyase activity appears to be critical for utilization of CS-B as a sole carbon source. The PMI2128 mutant also exhibited reduced growth on CS-C compared to the wild-type strain (Figure 4E), further suggesting a CS-type specificity for exolyase activity. It is also notable that the PMI2127/2128 mutant failed to grow under any condition except when supplied with glucose (Supplemental Figure 3E), confirming that these are the only enzymes the contribute to CS degradation in strain HI4320.

Since wild-type and the PMI2128 mutant both grew better on the unspecified CS mixture than any individual CS type, we next sought to determine if a 1:1 mixture of two CS types (CS-A and CS-C) could bolster growth (Figure 3F). Wild-type *P. mirabilis* had significantly better growth on the 1:1 mixture than either CS type alone, reaching a density approaching that of the glucose condition and in line with the chondroitin mixture condition (OD_600_>0.4, Figure 4F). The PMI2124 mutant had a similar growth rate as wild-type on the 1:1 mixture, but plateaued at a lower optical density suggesting that sulfatase activity is less critical when growing on a mixture of CS types but still required for optimal degradation and utilization as a carbon source. The PMI2127 mutant barely grew, confirming that endolyase activity is the most critical for CS utilization, while the PMI2128 mutant had a pronounced growth defect but achieved stationary phase at the same density as the PMI2124 mutant, indicating that exolyase activity also contributes to the ability to use CS as a carbon source.

### *P. mirabilis* chondroitin degradation contributes to infection

Considering that CS is the most abundant GAG in the urinary tract, we next sought to determine the contribution of CS degradation to *P. mirabilis* pathogenesis within the urinary tract (33, 34). To investigate the role of *P. mirabilis* chondroitinases during infection, we first utilized the murine CAUTI model in which female CBA/J mice were transurethrally infected with 1×10^5^ CFU/mouse of either wild-type *P. mirabilis* or each CS utilization mutant. Bacterial burden in the urine, bladder, kidneys, and spleen was determined at 48 hours post infection (Figure 5A). In the urine and bladder, the PMI2127 mutant displayed significantly reduced CFUs compared to the wild-type, while also showing a trend of reduced CFUs in the kidneys that was not statistically significant. The PMI2124 and PMI2128 mutants colonized all organs to a similar level as wild-type *P. mirabilis,* suggesting that sulfatase and exolyase activity are not required for CAUTI pathogenesis. However, the PMI2127/2128 double mutant showed a trend of reduced CFUs in the urine, significantly reduced CFUs in the bladder and kidneys, and a significant reduction in the overall incidence of bladder and kidney colonization compared to wild-type (Figure 5A). Thus, while endolyase activity alone strongly contributes to infection in this model, exolyase activity may provide some degree of functional redundancy either for circumventing the barrier of the urothelial GAG layer or supporting *P. mirabilis* growth in this environment.

**Figure 5.**
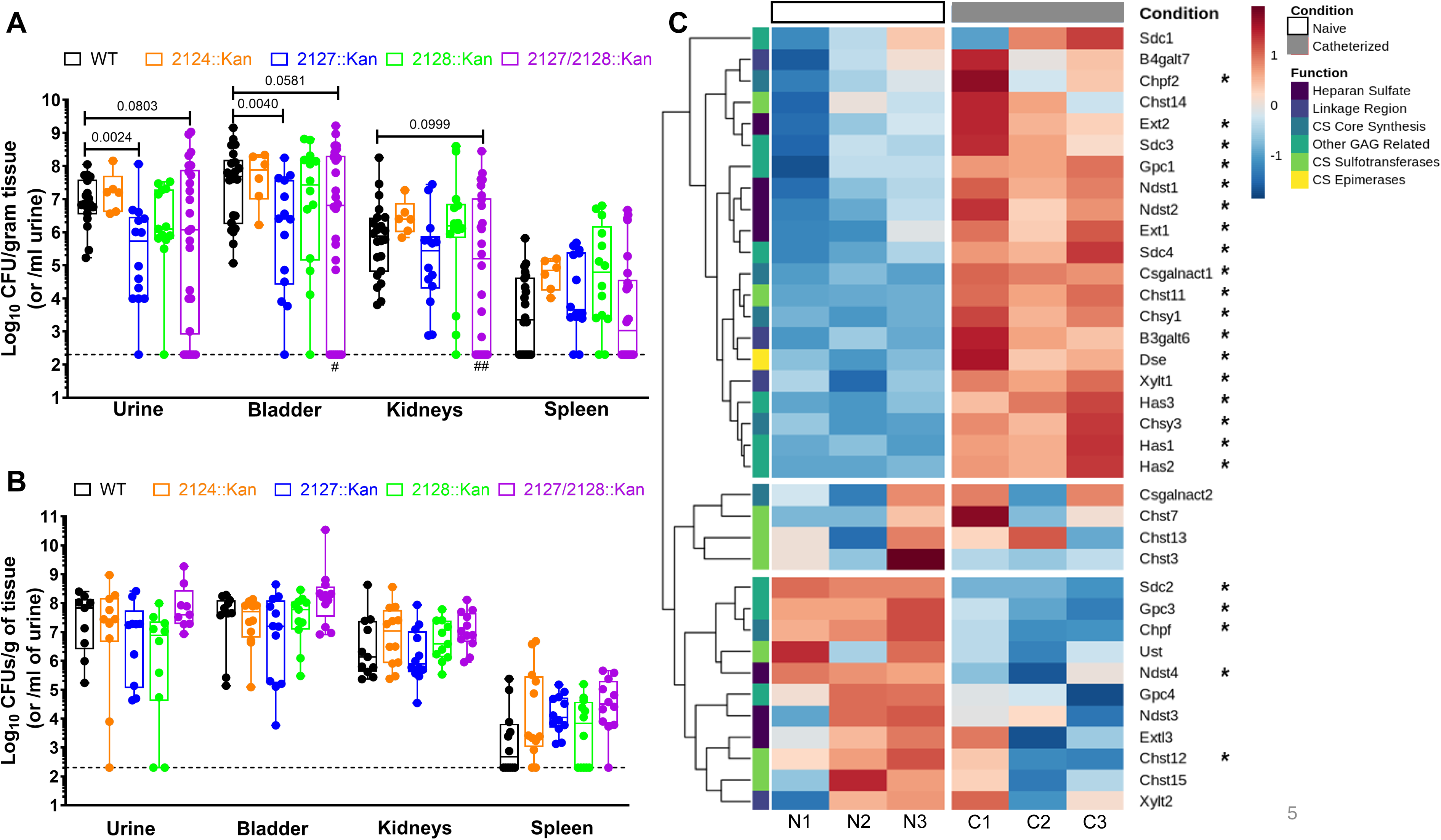
Contribution of *P. mirabilis* chondroitin degrading enzymes to infection. A) Female CBA/J mice were transurethrally inoculated with 10^5^ CFU of either wild-type *P. mirabilis* HI4320 (black circles), the PMI2124 mutant (orange circles), the PMI2127 mutant (blue circles), the PMI2128 mutant (green circles), or the PMI2127/2128 double mutant (purple circles), and a 4mm segment of silicone catheter tubing was placed in the bladder during inoculation to simulate CAUTI. After 48h, CFUs were determined in the urine, bladder, kidneys, and spleen. B) Female CBA/J mice were inoculated transurethrally with 10^7^ CFU of either wild-type *P. mirabilis* HI4320 (black circles), the PMI2124 mutant (orange circles), the PMI2127 mutant (blue circles), the PMI2128 mutant (green circles), or the PMI2127/2128 double mutant (purple circles) in a mouse model of ascending UTI (no catheter). Bacterial burden was quantified in the urine, bladder, kidneys, and spleen at 48 hours post infection. Groups were compared by one-way ANOVA of log-transformed CFUs. Dashed lines indicate limit of detection, gray numbers in panel A indicate percent of mice with CFUs above the limit of detection for each organ. #P<0.05, ##P<0.01 by Fischer’s exact test. C) Heat map displaying the impact of catheterization on expression of glycosaminoglycan-related genes in female B6 mice. Expression levels for three mice per condition are displayed, with gene function indicated on the left and asterisks indicating adjusted *P*<0.05 on the right.

There is also a GAG glycocalyx that extends into the vascular lumen, which participates in numerous functions including being a physical barrier to pathogens and mechanosensation (35). We therefore sought to determine if CS degradation activity by *P. mirabilis* contributes to systemic infection or kidney colonization during hematogenous spread. Bacteremia infection studies were conducted by inoculation female CBA/J mice 7 weeks of age via tail vein injection with 1×10^7^ CFU/mouse of either wild-type *P. mirabilis* or the PMI2127/2128 double mutant, since this mutant had the most robust defects in the CAUTI model. Animals were sacrificed 24 hours post-infection and bacterial counts in the liver, kidneys and spleen were determined (Supplemental Figure 4). In comparison to wild-type, the PMI2127/2128 double mutant had similar CFUs in all organs. However, mice infected with the double mutant had a lower overall incidence of liver colonization. Thus, CS lyase activity is not critical for the ability of *P. mirabilis* to survive within the bloodstream or infect the kidneys by hematogenous spread, but it may contribute to colonization of other secondary organs such as the liver.

While *P. mirabilis* is most well known as a cause of CAUTI, this species is also a common cause of complicated infection in non-catheterized individuals. To determine whether the contribution of chondroitin sulfate degradation to infection is specific to the catheterized urinary tract or a general feature of *P. mirabilis* UTI, we repeated our infection studies in a murine UTI model without simulating CAUTI. Female CBA/J mice were transurethrally inoculated with 1×10^7^ CFU/mouse of either wild-type *P. mirabilis* or the CS degradation mutants via a flexible pediatric venous catheter that was immediately removed after instillation of the inoculum, and bacterial burden in the urine, bladder, kidneys, and spleen was determined 48 hours post infection. Interestingly, all strains colonized to a similar level as wild-type *P. mirabilis* (Figure 5B), indicating that CS degradation only contributes to infection in the context of the catheterized bladder.

We hypothesized that the differential contribution of CS degradation to *P. mirabilis* colonization during CAUTI but not during UTI could be due to an altered GAG landscape in the presence of a catheter. To determine whether catheterization alters expression of genes involved in GAG synthesis or modification, we analyzed a publicly-available RNA-seq dataset from Rousseau et al. (36). In the referenced study, RNA was extracted from whole bladders of 6-8-week-old naïve female C57BL/6 mice 24 hours after implantation of a silicone catheter segment to examine the impact of catheterization on bladder gene expression. Raw sequencing reads from Rousseau et al. were downloaded from the NCBI sequence repository, trimmed, aligned to the mouse reference genome, and differential expression analyzed using DESeq2. A curated list of 35 GAG-related genes was then compiled based on the KEGG Glycosaminoglycan Biosynthesis pathway for targeted analysis of the impact of catheterization on expression of GAG-related genes.

Of the 35 GAG-related genes in KEGG, 18 (51%) were significantly increased in catheterized mice compared to naïve mice and 5 (14%) were significantly decreased (Figure 5C). The upregulated genes include 6 that are specific to CS synthesis or modification (*Csgalnact1, Chst11, Chys1*, *Chys3, Dse, Chpf2*) and 3 for HA (*Has1, Has2, Has3*). The five downregulated genes include 2 pertaining to the transfer of sulfate groups on CS (*Chst12*) and elongation of CS polymers (*Chpf*), with the other being involved in synthesis of proteoglycans and heparan modification. Thus, the presence of a catheter segment in the mouse bladder stimulates a transcriptional response that is likely to alter the GAG landscape of the bladder and therefore also the necessity for bacterial GAG-degrading enzymes during infection.

## Discussion

Glycosaminoglycans are found throughout the body and are thought to comprise an essential component of bladder barrier function (20, 37, 38). However, few studies have addressed the importance of the urothelial GAG layer during urinary tract infection or the interactions that occur between this layer and the uropathogens that frequently invade this body site. As a physical barrier and one of the first lines of defense from infection, it is important to understand how the GAG layer is degraded by uropathogens, such as *P. mirabilis.* In this study, we expanded on the initial observations made by Nguyen et al. 2022 (17) to demonstrate that *P. mirabilis* is a prolific degrader of the major urothelial GAG (chondroitin sulfate, CS) under numerous media conditions, and further define the contribution of three genes of interest to degradation and to infection in three different models. Critically, we determined that CS degradation only contributes to *P. mirabilis* infection in the catheterized urinary tract and provide evidence that this specificity is due to the impact of the catheter on GAG synthesis and modification pathways in bladder tissue. Collectively, our results highlight a new player mediating host-pathogen interactions in the catheterized bladder.

This study contributes to a growing body of work addressing uropathogen interactions with urinary GAGs and their role in pathogenesis. Earlier work had shown that *Proteus vulgaris*, a commensal gut species and occasional cause of CAUTI, was able to degrade chondroitin sulfate and lead to characterization of chondroitin sulfate endolyase and exolyase enzymes that are homologs to those examined in this study (39–41). However, this prior work focused on basic science characterizations of enzymatic activity and protein structure and had not been expanded to other pathogens nor placed in the context of infection. Through targeted mutagenesis we demonstrate that the predicted endolyase in *P. mirabilis* (PMI2127) is essential for GAG degradation activity, while exolyase activity via PMI2128 only contributes to utilization of a subset of CS types. Interestingly, the PMI2124 sulfatase was also essential for efficient GAG degradation by *P. mirabilis*. Thus, removal of the sulfate groups decorating the glycan sugars may be critical for *P. mirabilis* to properly recognize or access CS for degradation, as has been described for the ability of certain gut bacteria to degrade mucins (42).

The contribution of PMI2128 to growth with CS as a sole carbon source was initially unexpected considering that complete degradation of all CS substrates was achieved in the semi-quantitative GAG degradation assay. However, it is important to note that the GAG degradation assay only detects high molecular weight GAGs capable of binding BSA and precipitating upon acidification (17). Thus, it is possible that PMI2128 exolyase activity is sufficient to reduce CS to small enough fragments that no precipitation occurs in this assay without providing sufficient cleavage of disaccharides for use as a carbon source. The exact size and nature of the degradation products of PMI2127 and PMI2128 remains to be determined.

It is notable that the time frame of CS degradation by *P. mirabilis* differed under different growth conditions, occurring earlier in the more nutrient-limited urine than rich LB. Human urine is thought to predominantly provide bacteria with amino acids and small peptides as the primary nutrient sources (3, 43, 44). However, the presence of host GAGs may supply a more favorable carbon source to support rapid growth through a mechanism that is unneeded in LB until all other nutrients have been exhausted (45). Other GAG degrading species found in the gut, such as *B. thetaiotaomicron,* have been shown to utilize degraded GAGs for growth and disruption of GAGase activity resulted in significant growth lags in this species (32, 46). Thus, GAG degradation and utilization may contribute to bacterial growth in numerous sites within the body. The utilization of CS to sustain *P. mirabilis* growth likely occurs to varying degrees in different tissues within the host, dependent upon the nutrient conditions present during infection and possibly nutrient competition with other species. Further work is needed to elucidate the conditions under which *P. mirabilis* engages in GAG degradation, including identification of any metabolic conditions or signals that regulate expression and activity of GAGases.

Utilizing a murine CAUTI and bacteremia model, we demonstrated that disruption of CS degradation has a significant impact on bacterial counts during CAUTI but not within bacteremia. Interestingly, the PMI2124 mutant did not show a defect *in vivo*, indicating that the CS present within the mouse bladder may be already processed in such a way that sulfatase activity is redundant, whilst in the *in vitro* assay it was required. The lack of a defect *in vivo* was unexpected, as previous work demonstrated that sulfatase activity is needed in order for bacteria to access GAGs present within the gut for degradation (47). However, this discrepancy could be due to altered sulfonation patterns on GAGs within the catheterized bladder environment, or the activity of host enzymes may allow for more permissive access to GAGs in the bladder. Additionally, the lack of colonization defect for the PMI2124 mutant indicates that loss of degradation activity *in vitro* was not due to polar effects on the rest of the chondroitinase operon, despite the insufficiency of complementation *in vitro*.

While GAGs are present in both the bladder and arterial tissues (48, 49), it appears that *P. mirabilis* degradation of CS is tissue-specific and most important during bladder infection, but only in the context of catheterization. One explanation for the differential contribution of CS degradation to CAUTI and not UTI may be the host response induced by the catheter itself. Catheter insertion alone can induce tissue damage and inflammation, which in turn necessitates the host to engage in wound healing and remodeling, a process that heavily involves GAGs including chondroitin sulfate (50–55). Thus, it is possible that degradation of chondroitin sulfate becomes more important during CAUTI in order to deal with an altered GAG landscape due to tissue remodeling induced by the host. If so, this has implications for examining microbe-GAG interactions in a wide range of pathogens that are more commonly associated with CAUTI than uncomplicated UTI.

A caveat for our data is the focus on one urinary isolate of *P. mirabilis* to characterize GAG degradation capability. Even though >99% of 1,027 *P. mirabilis* urine isolate genomes contained all three chondroitin sulfate degrading genes, the degradation capabilities of each may be different than that seen in *P. mirabilis* HI4320. However, this study builds a framework wherein other clinical isolates of *P. mirabilis* can be tested to determine the heterogeneity of GAG degradation. Strain-dependent differences in GAG degradation have been seen in gut colonizers such as *Bacteroides,* where there are distinct groups of CS-HA degraders, CS only degraders, and non-degraders (46, 56).

Since the catheterized bladder appears to be an environment where GAGs are plentiful and catheterization can induce damage and a host response that likely increases GAG availability, GAG degradation may be an activity shared by numerous clinical strains of *P. mirabilis* as well as other species (36). As mentioned above, chondroitinase activity was first characterized in *P. vulgaris*, another frequent and persistent colonizer of the catheterized urinary tract (57, 58).

Thus, investigations into how GAG degradation impacts *P. vulgaris* infection are a natural next step. Additionally, while this work focused on mono species infections, the catheterized bladder environment is often polymicrobial and there is the potential for competition or cross-feeding interactions mediated by these GAGs to occur (58–60). Further work is therefore needed to address *P. mirabilis* GAG degradation in more complex, multi-species contexts.

## Methods

### Analysis of *P. mirabilis* genomes for presence of GAG genes

*Proteus mirabilis* genomes (n=1,027) annotated using Prokka v1.14.6 were screened for three virulence genes (PMI2124 sulfatase CAR44241, PMI2127 chondroitin ABC endolyase CAR44246, PMI2128 chondroitin ABC lyase CAR44248) (30, 61). These genes were chosen based on their established roles in *P. mirabilis* pathogenesis and availability of validated reference sequences from the HI4320 strain. Sequence validation was performed using DIAMOND v2.1.11 with stringent criteria (≥80% identity, ≥80% coverage, e-value ≤1e-10) (62). Annotation accuracy was assessed via sensitivity, specificity, and F1-score calculations.

### Bacterial Strains

*Proteus mirabilis* strain HI4320 was isolated from the urine of a long-term catheterized patient in a chronic care facility (63). All *P. mirabilis* mutants used in this study were generated by inserting a kanamycin resistance cassette into the gene of interest following the Sigma TargeTron group II intron protocol as previously described (64). Mutants were verified by selection on kanamycin and PCR. The complemented strains were generated by cloning the appropriate gene with ∼500bp flanking regions into pGen-Amp and verified by PCR. Empty vectors strains contain only the pGen-Amp plasmid. Strains are listed with their genetic modifications and antibiotic resistance in Table 2.

**Table 1.**
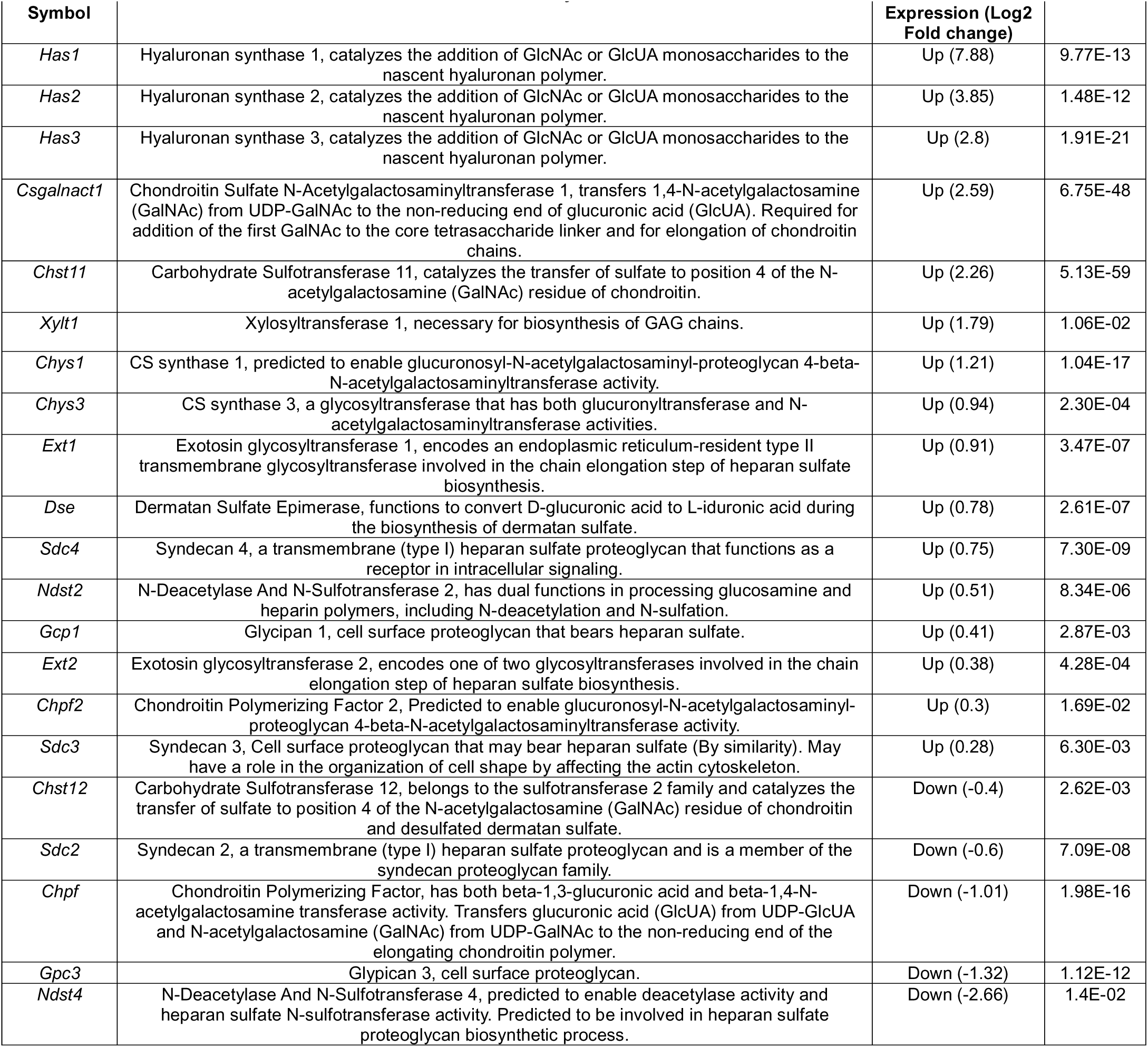
GAG associated genes of interest that are differentially expressed in catheterized vs naïve B6 mice.

| Symbol |  | Expression (Log2 Fold change) |  |
| --- | --- | --- | --- |
| *Has1* | Hyaluronan synthase 1, catalyzes the addition of GlcNAc or GlcUA monosaccharides to the nascent hyaluronan polymer. | Up (7.88) | 9.77E-13 |
| *Has2* | Hyaluronan synthase 2, catalyzes the addition of GlcNAc or GlcUA monosaccharides to the nascent hyaluronan polymer. | Up (3.85) | 1.48E-12 |
| *Has3* | Hyaluronan synthase 3, catalyzes the addition of GlcNAc or GlcUA monosaccharides to the nascent hyaluronan polymer. | Up (2.8) | 1.91E-21 |
| *Csgalnact1* | Chondroitin Sulfate N-Acetylgalactosaminyltransferase 1, transfers 1,4-N-acetylgalactosamine (GalNAc) from UDP-GalNAc to the non-reducing end of glucuronic acid (GlcUA). Required for addition of the first GalNAc to the core tetrasaccharide linker and for elongation of chondroitin chains. | Up (2.59) | 6.75E-48 |
| *Chst11* | Carbohydrate Sulfotransferase 11, catalyzes the transfer of sulfate to position 4 of the N-acetylgalactosamine (GalNAc) residue of chondroitin. | Up (2.26) | 5.13E-59 |
| *Xylt1* | Xylosyltransferase 1, necessary for biosynthesis of GAG chains. | Up (1.79) | 1.06E-02 |
| *Chys1* | CS synthase 1, predicted to enable glucuronosyl-N-acetylgalactosaminyl-proteoglycan 4-beta-N-acetylgalactosaminyltransferase activity. | Up (1.21) | 1.04E-17 |
| *Chys3* | CS synthase 3, a glycosyltransferase that has both glucuronyltransferase and N-acetylgalactosaminyltransferase activities. | Up (0.94) | 2.30E-04 |
| *Ext1* | Exotosin glycosyltransferase 1, encodes an endoplasmic reticulum-resident type II transmembrane glycosyltransferase involved in the chain elongation step of heparan sulfate biosynthesis. | Up (0.91) | 3.47E-07 |
| *Dse* | Dermatan Sulfate Epimerase, functions to convert D-glucuronic acid to L-iduronic acid during the biosynthesis of dermatan sulfate. | Up (0.78) | 2.61E-07 |
| *Sdc4* | Syndecan 4, a transmembrane (type I) heparan sulfate proteoglycan that functions as a receptor in intracellular signaling. | Up (0.75) | 7.30E-09 |
| *Ndst2* | N-Deacetylase And N-Sulfotransferase 2, has dual functions in processing glucosamine and heparin polymers, including N-deacetylation and N-sulfation. | Up (0.51) | 8.34E-06 |
| *Gcp1* | Glycipan 1, cell surface proteoglycan that bears heparan sulfate. | Up (0.41) | 2.87E-03 |
| *Ext2* | Exotosin glycosyltransferase 2, encodes one of two glycosyltransferases involved in the chain elongation step of heparan sulfate biosynthesis. | Up (0.38) | 4.28E-04 |
| *Chpf2* | Chondroitin Polymerizing Factor 2, Predicted to enable glucuronosyl-N-acetylgalactosaminyl-proteoglycan 4-beta-N-acetylgalactosaminyltransferase activity. | Up (0.3) | 1.69E-02 |
| *Sdc3* | Syndecan 3, Cell surface proteoglycan that may bear heparan sulfate (By similarity). May have a role in the organization of cell shape by affecting the actin cytoskeleton. | Up (0.28) | 6.30E-03 |
| *Chst12* | Carbohydrate Sulfotransferase 12, belongs to the sulfotransferase 2 family and catalyzes the transfer of sulfate to position 4 of the N-acetylgalactosamine (GalNAc) residue of chondroitin and desulfated dermatan sulfate. | Down (-0.4) | 2.62E-03 |
| *Sdc2* | Syndecan 2, a transmembrane (type I) heparan sulfate proteoglycan and is a member of the syndecan proteoglycan family. | Down (-0.6) | 7.09E-08 |
| *Chpf* | Chondroitin Polymerizing Factor, has both beta-1,3-glucuronic acid and beta-1,4-N-acetylgalactosamine transferase activity. Transfers glucuronic acid (GlcUA) from UDP-GlcUA and N-acetylgalactosamine (GalNAc) from UDP-GalNAc to the non-reducing end of the elongating chondroitin polymer. | Down (-1.01) | 1.98E-16 |
| *Gpc3* | Glypican 3, cell surface proteoglycan. | Down (-1.32) | 1.12E-12 |
| *Ndst4* | N-Deacetylase And N-Sulfotransferase 4, predicted to enable deacetylase activity and heparan sulfate N-sulfotransferase activity. Predicted to be involved in heparan sulfate proteoglycan biosynthetic process. | Down (-2.66) | 1.4E-02 |

**Table 2.**
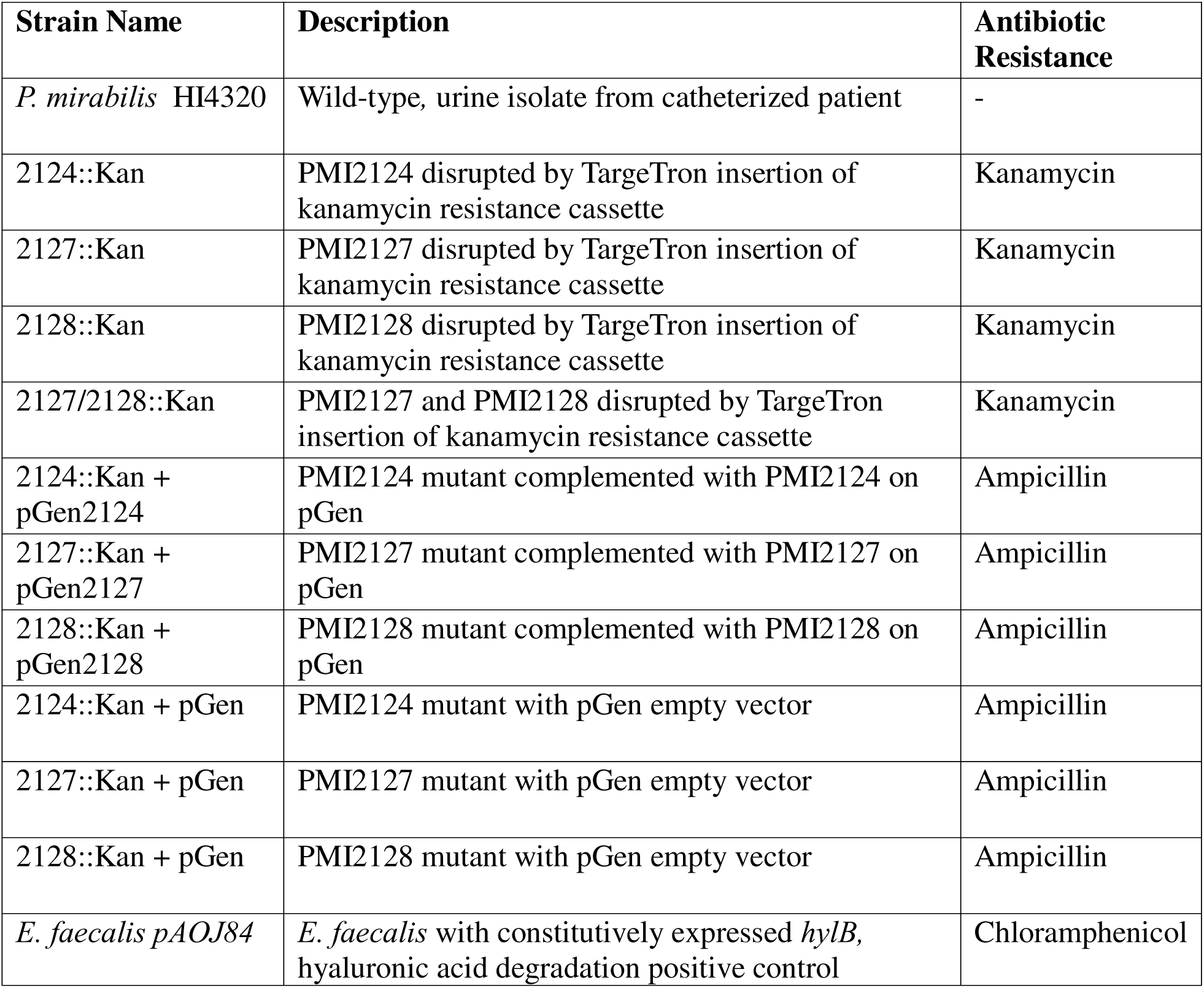
Bacterial strains used in this study. Wild-type *P. mirabilis* HI4320, chondroitin sulfate mutants, and complemented strains are listed with their modifications and antibiotic resistance profiles.

### Growth and Media Conditions

*P. mirabilis* was cultured at 37°C with shaking at 225 RPM in 5 mL of low-salt LB broth (10 g/L tryptone, 5 g/L yeast extract, 0.1g/L NaCl) or on plates solidified with 1.5% agar. Mutant strains were cultured with 50 µg/mL kanamycin, while complemented strains and those with empty vectors were cultured with the addition of 100 µg/mL of ampicillin to maintain plasmid selection. *Proteus mirabilis* minimal salts medium (PMSM) lacking a carbon source was used in experiments requiring defined growth medium: 10.5 g/L K_2_HPO_4_, 4.5 g/L KH_2_PO_4_, 1 g/L (NH_4_)_2_SO_4_, 15 g/L agar, supplemented with 0.002% nicotinic acid, and 1 mM MgSO_4_. The no-carbon PMSM was then supplemented with 2.5 mg/mL of either chondroitin sulfate or glucose. Filter sterilized pooled human urine from at least 20 de-identified female donors was purchased from Cone Bioproducts (Sequin, TX) and stored at −20C until use. Urine aliquots were then diluted 1:1 with 0.9% saline to generate a dilute urine medium suitable for *P. mirabilis* growth without excessive precipitation from urease activity (65). Where indicated, media were supplemented with the following glycosaminoglycans (GAGs): chondroitin sulfate (Thermo, Cat# J60341.14), chondroitin sulfate A (Thermo, Cat# C9819-5G), chondroitin sulfate B (Thermo, Cat# C3788-100MG), chondroitin sulfate C (AA blocks, Cat# AA00ILKC), hyaluronic acid (Sigma, Cat#53747), and heparan sulfate (Sigma, Cat# H3149). All GAGs were prepared as 5 mg/mL or 10 mg/mL stock solutions in Milli-Q water and filter sterilized using a 0.2µm filter.

### Characterization of GAGase activity

GAGase activity was determined as described in Nyugen et al. 2022 (66). In brief, overnight cultures were washed with phosphate buffered saline (PBS), adjusted to ∼10^7^ CFUs/mL in the desired media supplemented with 2.5 mg/mL final concentration of the desired GAG, and 200 µL were aliquoted into 96-well plates and incubated for up to 48 hours at 37 °C. After incubation, plates were centrifuged at 3000 x g for 10 minutes to pellet the bacteria and supernatants were collected and transferred to another 96-well plate. Serial two-fold dilutions were performed in 1x PBS, after which 90 µL of diluted supernatant was transferred to a 96-well plate, bovine serum albumin (BSA) (Fischer, Cat# BP9700100) was added to a final concentration of 1% to each well, and the OD_600_ measurements of each well were taken (pre acetic acid value) with a BioTek Synergy H1 plate reader. Next, 40 µL of 2M acetic acid (Fischer, Cat#A38C-212) was added to each well and stirred using the pipette tip and the OD_600_ measurements were taken again (post acetic acid values). The percentage of GAGs remaining were calculated using the following formula:

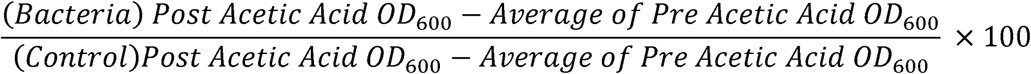

### Growth curves

Overnight cultures were washed with phosphate buffered saline (PBS), adjusted to ∼10^7^ CFUs/mL in *Proteus mirabilis* minimal salts media (PMSM) supplemented with 2.5 mg/mL final concentration of the desired GAG, glucose, or no carbon source, and 200 µL were aliquoted into 96-well plates. Cultures were incubated for 48 hours at 37 °C in the BioTek BioSpa 8 incubator, with OD_600_ readings taken every 15 minutes. Plates were orbitally shaken for 30 seconds prior to each OD reading for aeration.

### RT-qPCR of *P. mirabilis* chondroitinases

Samples for RNA extraction were prepared as described here (67). In brief, samples were prepared by inoculating 5 mL of LB with or without 2.5 mg/mL of chondroitin sulfate mix with the bacteria of interest and collecting pellets at 12, 18, or 24 hours. At each time point, the cultures were pelleted by spinning for 20 minutes at 8000 rcf, the supernatant was removed and the pellet was resuspended in 50 µL of RLT Buffer with BME (Qiagen RNeasy Mini Kit, Cat# 74104, Germany). Next, 1.5 mL safe-lock microfuge tubes (Eppendorf, Hamburg, Germany) were filled half way with 0.5 mm glass disruption beads (RPI Research Products International, Cat# SI-BG05), the resuspended pellet was then added, and samples were homogenized using a Bullet Blender Gold (Next Advance, Troy, NY, USA) at max speed for 5 minutes. The homogenate was collected and centrifuged at 21,000 rcf for 10 minutes at 4 °C, supernatant was removed, and 1 volume of 70% ethanol was added and vortexed to mix. The sample was then transferred to an RNeasy mini spin column (Qiagen) and spun through twice to ensure maximum binding. DNA digestion was performed on column using the DNAse digestion kit (Qiagen, Cat#79254, Germany). After digestion and washing the column, a second DNase digestion was performed off column using DNase Treatment & Removal Kit (Invitrogen, Cat# AM1906). cDNA was synthesized using the iScript cDNA Synthesis Kit (BioRad, Cat#1708890), and RT-qPCR was performed as previously described (68) using qPCRBio SyGreen Blue Mix Lo-Rox (PCR Biosystems, Cat# PB20.15-01) with a BioRad CFX-Connect Real Time system. Data were normalized to *recA* as the reference genes and analyzed via the Pfaffl method to determine relative fold gene expression of the samples while taking into account primer efficiencies (69).

### Murine CAUTI Model

Female CBA/J mice aged 6-8 weeks (Jackson Laboratory) were anesthetized with a weight appropriate dose of ketamine/xylazine (80-120mg/kg ketamine and 5-10 mg/kg xylazine) via IP injection and transurethrally inoculated with 50 µL of 2×10^6^ CFU/mL (1×10^5^ CFU/mouse) of wild-type *P. mirabilis,* 2124::Kan, 2127::Kan, 2128::Kan, or 2127/2128::Kan. A 4 mm segment of sterile silicone tubing (0.64 mm O.D., 0.30 mm I.D., Braintree Scientific Inc.) was advanced into the bladder during inoculation to be retained for the duration of the study as described previously (70, 71). After 48 hours, urine was collected, bladders, kidneys, and spleens were harvested and placed into 5 mL Eppendorf tubes containing 1 mL 1x PBS and 500 µL of 3.2mm stainless steel beads. Tissues were homogenized using a Bullet Blender 5 Gold (Next Advance, Speed 8, 4 minutes). Bladders were treated to two cycles to ensure full homogenization. Tissue homogenates were serially diluted and plated onto appropriate agar using an EddyJet 2 spiral plater (Neutec Group) for determination of CFUs using a ProtoCOL 3 automated colony counter (Synbiosis).

### Murine Bacteremia Model

Infections were carried out as described previously (72, 73). In brief, female CBA/J mice aged 7 weeks were inoculated via tail vein injection with 50 µL of 2×10^8^ CFU/mL (1×10^7^ CFU/mouse) of either wild-type *P. mirabilis* or the 2127/2128::Kan mutant. Mice were sacrificed 24 hours post infection and bacterial counts in the liver, kidneys, and spleen were quantified.

### Murine UTI Model

Infections were carried out as described previously (74). In brief, female CBA/J mice were inoculated transurethrally via a flexible pediatric venous catheter (Cat# 381512, Insyte^TM^ Autoguard^TM^ Winged, 0.7 x 19 mm) with 50 µL of 2×10^8^ CFU/mL (1×10^7^ CFU/mouse) of wild-type *P. mirabilis,* 2124::Kan, 2127::Kan, 2128::Kan, or 2127/2128::Kan mutant. Bacterial burden was quantified in the urine, bladder, kidneys, and spleen at 48 hours post infection as described above.

### Analysis of existing RNA-Seq dataset

RNA sequencing data were obtained from the NCBI Sequence Read Archive (BioProject PRJNA335539). The selected dataset comprised six samples from the study by Rousseau et al.: three samples from naive female C57BL/6 mouse bladders and three from bladders 24 hours post-catheterization (36). Raw sequencing reads were downloaded using the SRA Toolkit v3.0.5 and converted to paired-end FASTQ format. Read quality control and adapter trimming were performed using Trimmomatic v0.39 (75). The trimmed reads were aligned to the mouse reference genome (mm10/GRCm38) using HISAT2 v2.2.1 with default parameters (76). SAM files were converted to BAM format, coordinate-sorted, and indexed using SAMtools v1.16.1 (77). Gene-level quantification was performed using featureCounts v2.0.4 from the Subread package (78). Differential expression analysis was conducted using the DESeq2 package v1.36.0 in R (79). Next, a targeted analysis was performed on genes involved in chondroitin sulfate (CS) and other glycosaminoglycan (GAG) biosynthesis and modification. A curated list of 35 GAG-related genes was compiled based on the KEGG Glycosaminoglycan Biosynthesis pathway (mmu00532). Differential expression results for these genes were extracted from the full analysis for focused interpretation. Complete analysis documentation is available at https://github.com/Deka-nam/rna-seq/tree/main/GAG_project.

### Statistical analysis

All analyses were performed using GraphPad Prism, version 10.4 (GraphPad Software) with a 95% confidence interval.

## Acknowledgements

This work was supported by the National Institute of Health through award F32AI183755 to BH, and awards R01DK136875 and R01DK140371-01A1 to CEA.

---

**Supplemental Figure 1.** Expression levels of chondroitin sulfate degradation genes in *P. mirabilis.* (A) Relative expression of PMI2124, PMI2127, and PMI2128 in wild-type *P. mirabilis* grown in LB with 2.5 mg/mL of chondroitin sulfate at 12, 18 and 24 hours compared to LB without supplementation at 12 hours of growth. (B-D) Fold change in expression of B) PMI2124, C) PMI2127 and D) PMI2128 in mutant strains with empty vectors or complementation vectors relative to wild-type *P. mirabilis* HI4320 with an empty vector. All strains were grown in LB with 2.5 mg/mL of chondroitin sulfate mix and 100 µg/ml ampicillin to maintain plasmid selection. For all experiments, data were normalized to *recA* as the reference gene and fold change in expression was determined via the Pfaffl method accounting for primer efficiencies. Data represent mean ± SD for three independent experiments with two technical replicates each.

**Supplemental Figure 2.** *P. mirabilis* does not degrade hyaluronic acid or heparan sulfate. A) Percent hyaluronic acid remaining after 48hrs of growth of *P. mirabilis* HI4320 or indicated mutant strains in LB media. An *E. faecalis* strain constitutively expressing hyaluronidase (*hylB*) was used as a positive control for hyaluronic acid degradation. B) Percent heparan sulfate remaining after 48hrs of growth of *P. mirabilis* HI4320 or indicated mutant strains in LB media. Data represent mean ± SD of at least three independent experiments.

**Supplemental Figure 3.** *P. mirabilis* 48-hour growth curves in PMSM. Wild-type *P. mirabilis* HI4320 (A), the PMI2125 mutant (B), PMI2127 mutant (C), PMI2128 mutant (D), and a PMI2127/2128 double mutant (E) were grown in PMSM with either no carbon source (No C) or supplemented with 2.5 mg/mL chondroitin sulfate mix (CS Mix), chondroitin sulfate A (CS-A), chondroitin sulfate B (CS-B), chondroitin sulfate C (CS-C), or 2.5 mg/mL glucose (Gluc). Growth was measured by OD600 readings every 15 minutes. Data represent mean ± SEM of at least three independent experiments.

**Supplemental Figure 4.** Contribution of *P. mirabilis* chondroitin degradation to bacteremia. Female CBA/J mice were inoculated via tail vein injection of 10^7^ CFUs of either wild-type *P. mirabilis* HI4320 (black circles) or the PMI2127/2128 double mutant (purple circles). After 24hr, CFUs were determined in the liver, kidneys, and spleen to examine secondary organ colonization during bacteremia. Groups were compared by one-way ANOVA of log-transformed CFUs. Dashed lines indicate limit of detection, gray numbers in panels A and B indicate percent of mice with CFUs above the limit of detection for each organ. #P<0.05, ##P<0.01 by Fischer’s exact test.

**Supplemental Table 1.** RT-qPCR primers and primers used to generate, complement and verify mutants used in this study.

