## Supplemental Figures and Table for "Chondroitin sulfate degradation bolsters *Proteus mirabilis* growth and colonization of the catheterized urinary tract"

Supplement 1

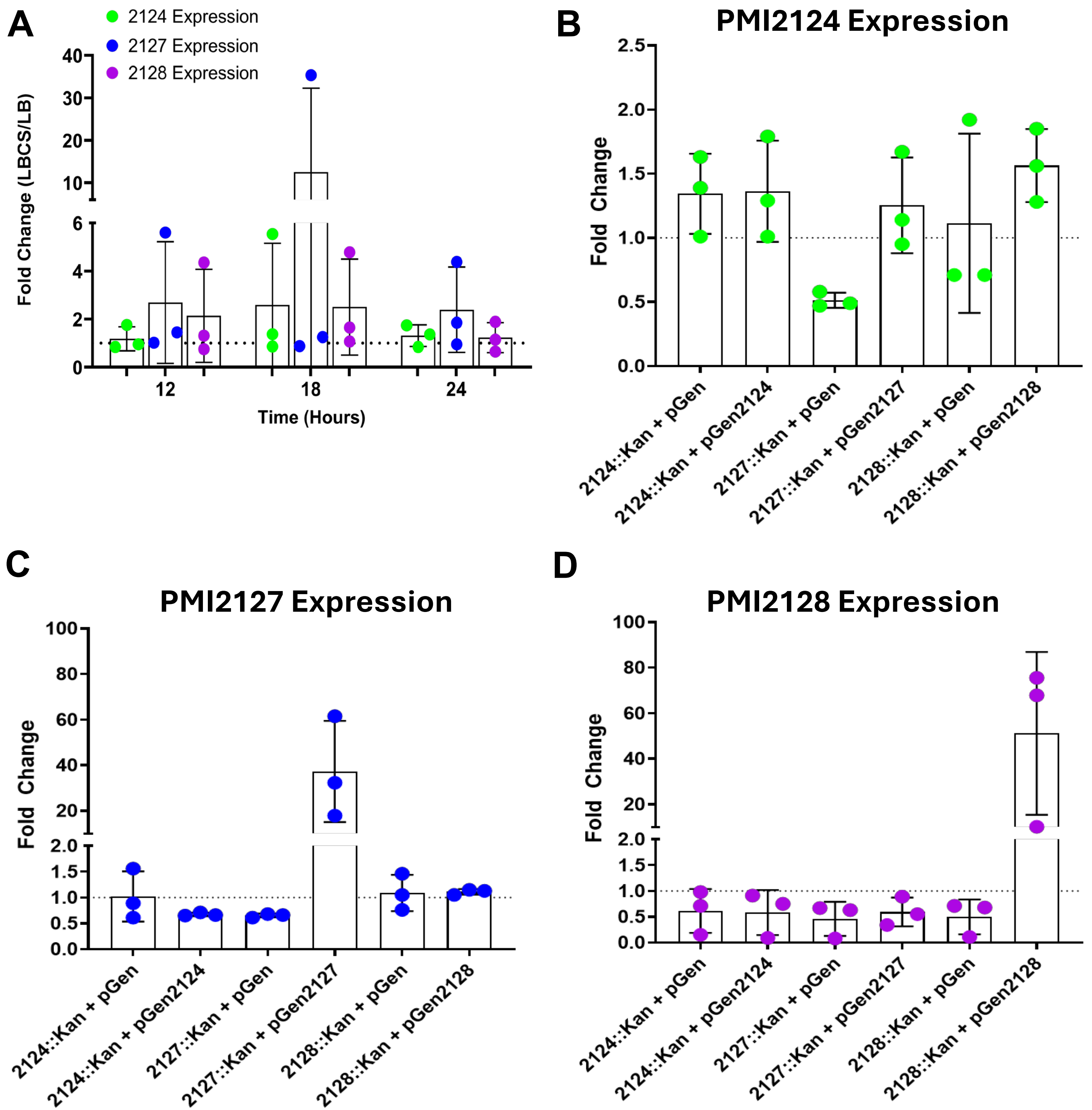

Supplement 2

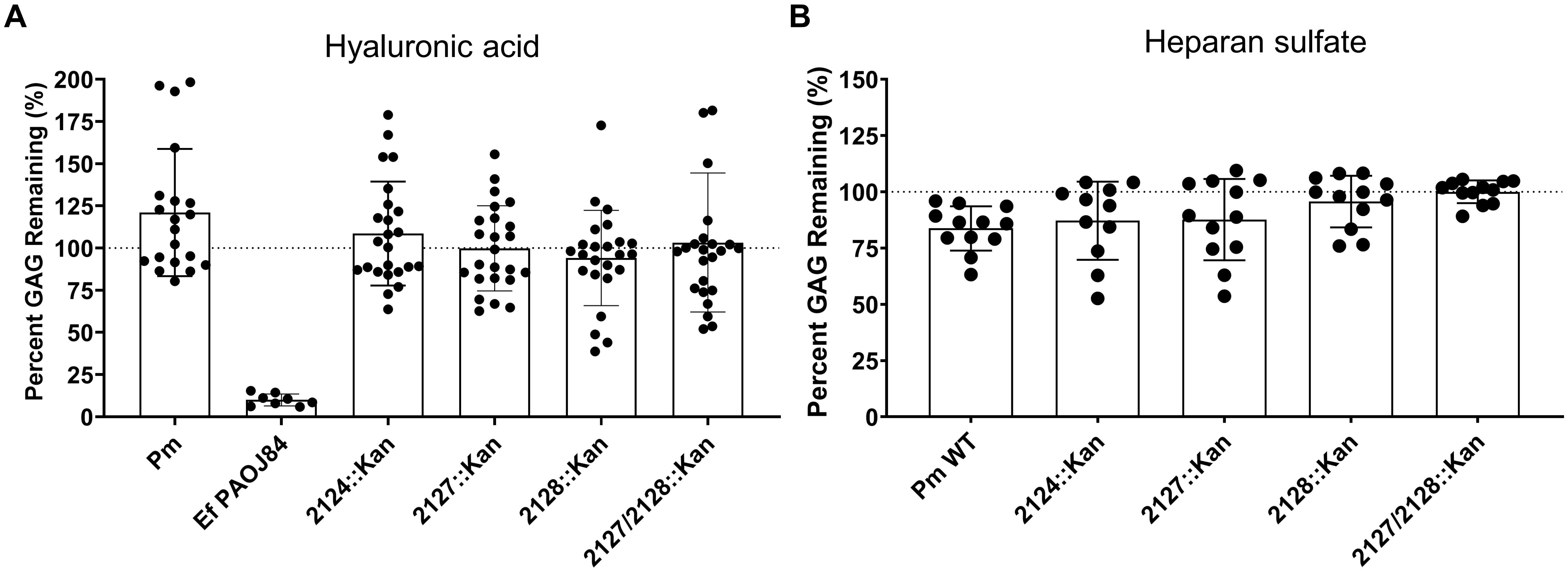

Supplemental 3

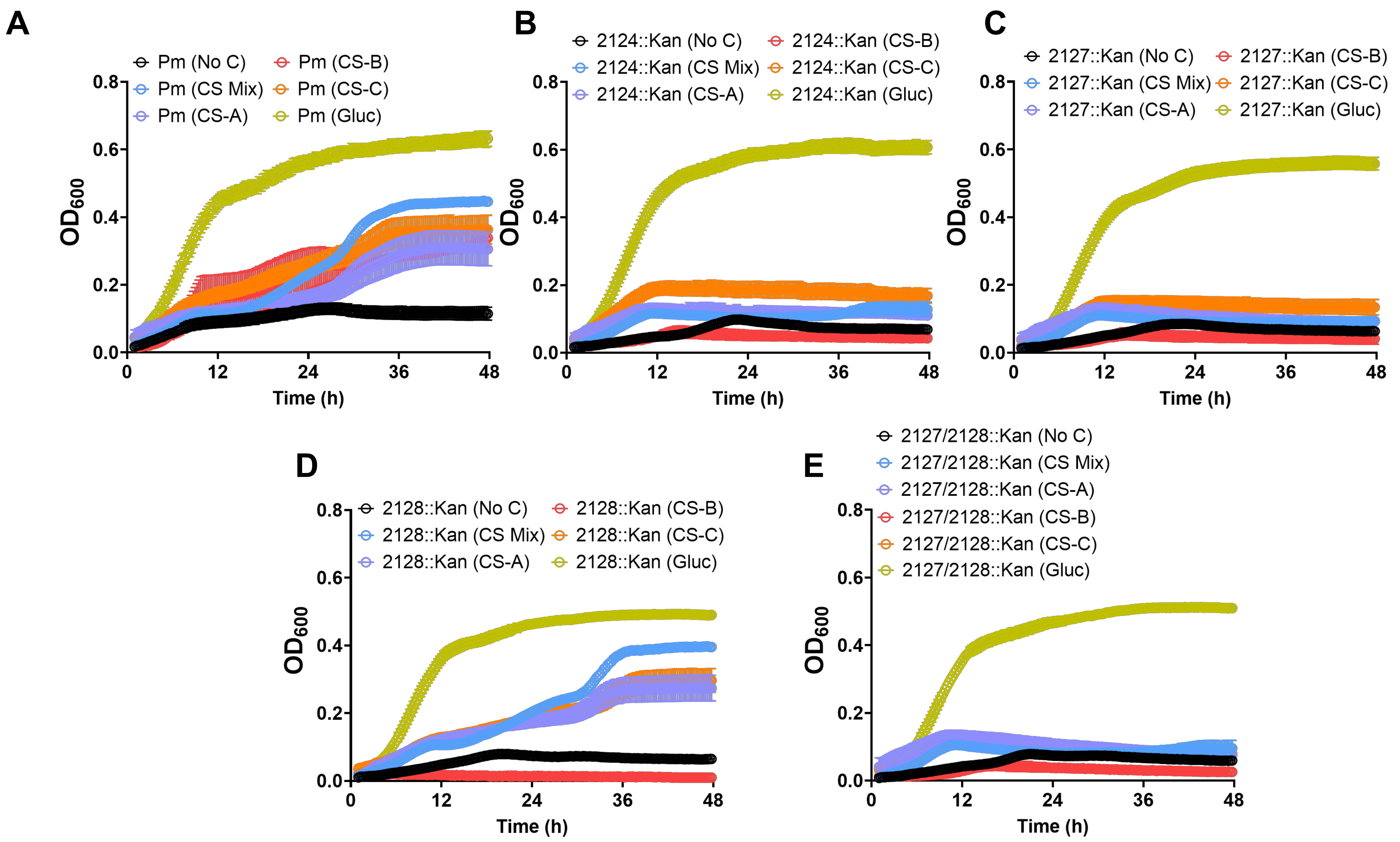


2.5 mg/mL final concentration of CS/gluc in PMSM with no other C source

Supplemental 4

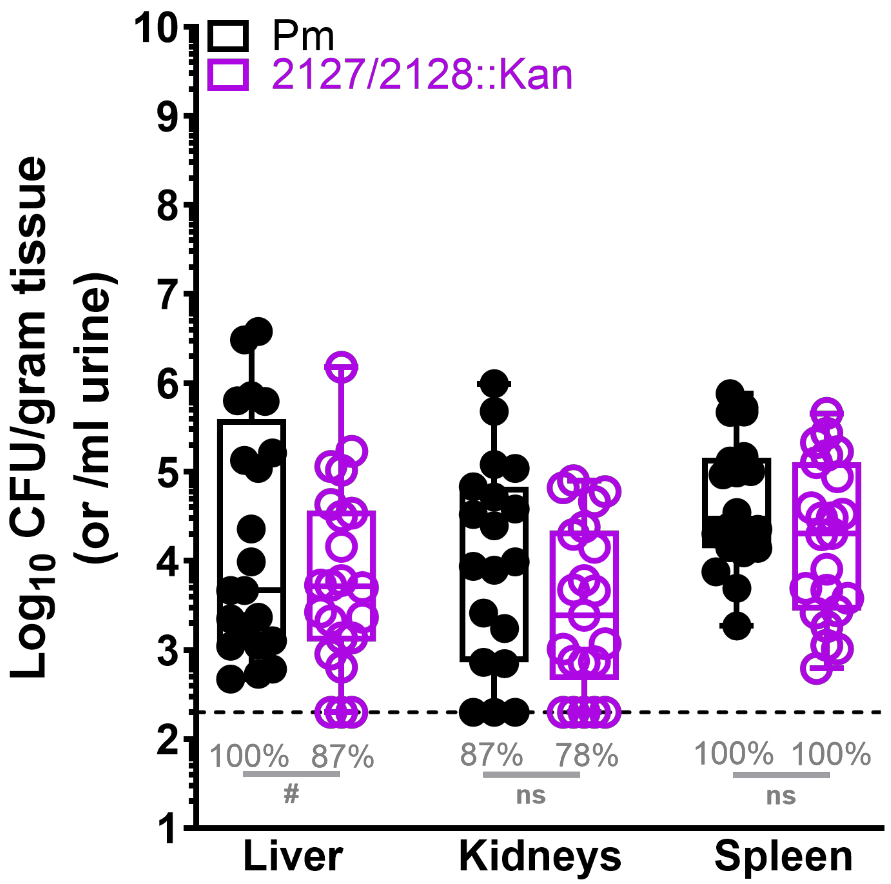

**Supplemental Table 1.** Primers used for RT-qPCR, mutant generation, verification, and complementation.

| Primer Name | Forward Primer | Reverse Primer |
|---|---|---|
| **RT-qPCR Primers** | | |
| recA (Housekeeping gene) | TCCGTGGCAGCATTAACA | TACGCATAGCTTGGCTCATC |
| rpoA (Housekeeping gene) | GCAAATCTGGCATTGGCCCTGTTA | TAGGGCGCTCATCTTCTTCCGAAT |
| PMI2127 RT-qPCR | CTGGCGTGCTATTGGTATCT | CTGTCAACATTCGCACCTAAAG |
| PMI2128 RT-qPCR | GCTAAAGGGCAAACGGTAGA | GGGCTTTACCGTGAGAGATAAC |
| **Mutagenesis, Complementation, and PCR Verification Primers** | | |
| PMI2124 Verification | GTTTCTACACCTGCTCTTGCTACTTCAGGT | AGACCAGCATTTTGCGTAAG |
| PMI2124 IBS | AAAAAAGCTTATAATTATCCTTACTCCACATAGGCGTGCGCCCAGATAGGGTG | |
| PMI2124 IBSd | CAGATTGTACAAATGTGGTGATAACAGATAAGTCCATGCTCTTAACTTACCTTTCTTTGT | |
| PMI2124 EBS2 | TGAACGCAAGTTTCTAATTTCGGTTTGGAGTCGATAGAGGAAAGTGTCT | |
| PMI2124 Comp | CGCTATTAACCGATCTTTACCAATA | TAAAAGTCGATAGTTAAACGGCTAC |
| PMI2127 Verification | AACAGCTAAAACAGATGTACCTCTT | CTATTCCAATCCCAGCCTTCTT |
| PMI2127 IBS | AAAAAAGCTTATAATTATCCTTACTAAACGAAGCTGTGCGCCCAGATAGGGTG | |
| PMI2127 IBSd | CAGATTGTACAAATGTGGTGATAACAGATAAGTCNNNNNNNNTAACTTACCTTTCTTTGT | |
| PMI2127 EBS2 | TGAACGCAAGTTTCTAATTTCGGTTTGAACGCAAGTTTCTAATTTCGGTTTTTAGTCGATAGAGGAAAGTGTCTTCGATAGAGGAAAGTGTCT | |
| PMI2127 Comp | AAGCTTGGTGTGCAATTAGGCTATCA | GGATCCGAGCCTTTCATATCACGAAAAG |
| PMI2128 Verification | TAGGCACTCGTTACGTTCTTG | CGTAGGATGTCTGCCACTTAAT |
| PMI2128 IBS | AAAAAAGCTTATAATTATCCTTAATCGTCGATGCAGTGCGCCCAGATAGGGTG | |
| PMI2128 IBSd | CAGATTGTACAAATGTGGTGATAACAGATAAGTCGATGCAAATAACTTACCTTTCTTTGT | |
| PMI2128 EBS2 | TGAACGCAAGTTTCTAATTTCGGTTACGATTCGATAGAGGAAAGTGTCT | |
| PMI2128 Comp | GAATTCACATGGTTTTCTTAGATGCAAC | GGATCCTCGAATTAGGAAATTGTGGGG |
